# The importance of selected markers of inflammation and blood-brain barrier damage for short-term post-stroke prognosis

**DOI:** 10.1101/503953

**Authors:** Anetta Lasek-Bal, Halina Jędrzejowska-Szypułka, Sebastian Student, Aldona Warsz-Wianecka, Katarzyna Zaręba, Przemysław Puz, Wiesław Bal, Katarzyna Pawletko, Joanna Lewin-Kowalik

**Author notes:** Address for correspondence: Anetta Lasek-Bal, MD, PhD, Department of Neurology, School of Health Sciences, Medical University of Silesia in Katowice, 40-735 Katowice, Ziołowa str. 45/47, Poland.

## Background

Stroke causes an immediate local immuno-inflammatory reaction. It is characterized by a strong activation of microglia, astrocytes and vascular endothelial cells; an inflow of inflammatory cells (primarily neutrophils, then macrophages and monocytes); an activation of adhesive molecules and the release of cytokines both from activated cells and the endothelium. Following brain ischemia, a non-specific response is triggered, resulting in the elimination of dead cells from the damage zone and in the protection of surviving neurons from excitotoxicity. In a later stage, specific response mechanisms are activated; these are significant for the regeneration and neuroplasticity [1].

The mechanism of post-stroke immune activation is poorly understood. Studies on experimental stroke suggest that cytokines modulate the process of neuronal damage during ischemic stroke. Although TNF and interleukin 1, 2 and 6 are the most frequently studied cytokines in stroke patients, the conclusions are ambiguous and inconsistent with the results of experimental stroke studies [2–4]. Increased blood concentrations of NSE and/or S100B after stroke indicate that the BBB has been damaged; they are associated with a worse neurological status. Still little is known about the importance of progranulin and other markers of post-stroke prognosis.

The aims of this project were: to assess the blood serum concentration of the selected markers of inflammation, of the BBB and nervous tissue damage, and of the coagulation system; we also wanted to assess the markers of neuro and angiogenesis in patients on the first day of ischemic stroke, and the mutual correlations between these marker levels. An additional aim was to assess the importance of the above parameters for the patient’s neurological state on the first day, and their functional status on the 30th day of stroke.

## Methods

Our prospective study conducted in 2016–2017 included patients with first-in-life stroke, as manifested clinically and identified according to WHO clinical criteria, who had an acute ischemic lesion of the brain visualized in computed tomography and/or magnetic resonance imaging of the head. The other main inclusion criteria were: the length of time from the onset of stroke symptoms to hospital admission ≤ 24 hours, and pre-stroke status according to modified Rankin Scale (mRS) ≤ 1 point.

All patients included in the study were analyzed according to:

- their age at first-in-life stroke;
- the presence of comorbidities, such as atrial fibrillation (AF), arterial hypertension (AH), coronary heart disease (CHD), diabetes mellitus (DM), lipid disorders (LD), > 70% atherosclerotic carotid artery stenosis (CAS, ipsilaterally to the acute ischemic brain lesion);
- their neurological status on the first day of stroke, evaluated on the NIHSS (National Institute of Health Stroke Scale); [5]
- plasma concentration of the following markers on the first day of stroke: interleukins IL-2 and IL-6; calcium-binding protein B, tumor necrosis factor alfa, progranulin, neuron-specific enolase, urokinase-type plasminogen activator (uPA), vascular endothelial growth factor (VEGF); brain-derived neurotrophic factor (BDNF), C reactive protein (CRP), leucocyte and platelet (PLT) counts;
- their functional status at 30 days following stroke as per the mRS scale. [6]

The diagnosis of AH was consistent with the recommendations of the European Society of Cardiology (ESC); DM was diagnosed according to the criteria of the Diabetes Association; dyslipidemia was defined according to the ESC recommendations (Guidelines for the Management of Dyslipidemias). [7–9]

The degrees of stenosis of the common carotid artery and/or internal carotid artery were assessed according to the NASCET criteria. [10]

Blood BDNF, Il-2, IL-6, S100B, TNF, GRN, NSE, uPA, VEGF concentrations on day 1 of stroke were assessed according to procedure as follows: 7 ml of EDTA blood was drawn and plasma was obtained after 15 minutes of centrifugation (1500 rpm), then the plasma was frozen at –80°C. The concentrations of above listed markers were measured using the ELISA method (Magnetic Luminex Assay R&D Systems). The assessments were performed at the Department of Physiology, Medical University of Silesia in Katowice.

Patients were categorized into three subgroups based on the NIHSS score (A ≤ 4, B 5–12 and C > 12). In each of these subgroups the mean concentrations of the examined substances were assessed and comparisons were made between them

Patients were categorized into three subgroups based on the mRS score (D ≤ 2 and E = 3, F > 3). In each of these groups formed the mean concentrations of the examined substances were assessed and comparisons were made between those subgroups.

A subgroup was formed of patients with symptomatic atherosclerosis of the coronary arteries (CHD and/or MI) and/or carotid arteries (>70% stenosis) / cerebral arteries (>50%) and ones with atherogenic stroke/TIA in the last 6 months, for which a profile of test concentrations was assessed and compared with the parameters in other patients.

A subgroup was formed of ‘the best clinical benefits’ (BB) patients; it consisted of those with scores > 10 on the NIHSS scale (the first day) who scored ≤2 on the mRS scale on the 30th day following the onset of stroke. For that subgroup, the profile of test concentrations was assessed and compared with the parameters in other patients.

Univariate analyses for differences between male and female groups for binary variables AH, DM, AF, LD, CHD, CAS, AT, therapy were performed using the Pearson’s Chi-squared test. Continuous variables were compared between male and female patients using Welch’s ANOVA or Kruskal-Wallis testing, as appropriate. We also analysed NIHSS and mRS grouped into 3-category ordinal scale. In this case for all not censored variables including WBC, PLT, CRP and cytokines we use a one-way ANOVA or a Kruskal-Wallis test. For censored data with values below the detection limit (LOD) we use the maximum likelihood estimation procedure or a Peto test accordingly. For all significant test results we performed post-hoc tests on pair-wise comparisons. The variable correlation plot was determined using the pearson correlation coefficient.

Multivariable models were built by using logistic regression and binomial GLM for dichotomous outcomes (eg, best benefits group) and ordinal logistic regression for ordinal outcomes (NIHSS, mRS). The model variable selection procedures included automatic selection (stepwise, forward and backward) based on AIC- and BIC criterion. For evaluating the accuracy of the model predictions the “leave one out” procedure and the area under the ROC curves (AUC) or multiclass AUC estimator was used. All statistical analyses were performed using R version 3.4.2.

The study was accepted by the Ethics Committee of the Silesian Medical University of Silesia in Katowice.

## Results

The study included 138 patients on the first day of ischemic stroke hospitalized in the Department of Neurology, Medical University of Silesia in Katowice: 75 patients were women, which accounted for 53.96% of all subjects. Mean age of the study subjects: 73.11 ± 11.48 [36–103]. The mean age of women was significantly higher; the neurological status on the first day of stroke and the functional status on the 30th day were significantly more severe than those in men. Mean PLT count and the incidence of DM were significantly higher among women. In the comparative analysis, the differences between other parameters during inclusion were not statistically significant.

The clinical characteristic of the patients is presented in Table 1 (Tab. 1).

**Table 1.** Characteristics of the study participants

| Parameter | All | Female | Male | p-value |
| --- | --- | --- | --- | --- |
| Age (years), mean± SD<br>[min, max] | 73.11± 11.48<br>[36-103] | 75.42± 10.92<br>[43-103] | 70.40± 11.71<br>[36-92] | 0.0101* |
| NIHSS, median (IQR)<br>[min, max] | 3 (6)<br>[1-18] | 5 (8)<br>[0-18] | 2 (3.5)<br>[0-15] | 0.0007*** |
| mRS, median (IQR)<br>[min, max] | 2 (3)<br>[0-6] | 3 (3)<br>[0-6] | 1 (2)<br>[0-5] | 0.0314* |
| ST, n, (%)<br>(A/S/C/O/D) | 19/45/40/33/2<br>(14/32/29/24/1) | 10/29/20/15/1<br>(13/39/27/20/1) | 9/16/20/18/1<br>(14/25/31/28/2) | 0.5582 |
| WBC, mean± SD<br>[min, max] | 8.52±3.09<br>[3.02-18.94] | 8.53± 3.06<br>[3.43-17.32] | 8.50± 3.15<br>[3.02-18.94] | 0.9545 |
| PLT, mean± SD<br>[min, max] | 233.67±69.11<br>[76-542] | 255.12±65.05<br>[121-542] | 211.71± 67.84<br>[76-435] | 0.0005*** |
| CRP, median (IQR)<br>[min, max] | 5 (17.92)<br>[0.5-218] | 5.05 (17.4)<br>[0.5-160] | 5 (19.17)<br>[5-218] | 0.9389 |
| AH, n (%) | 127 (91) | 68 (91) | 59 (92) | 0.7681 |
| DM, n (%) | 49 (35) | 26 (35) | 23 (36) | 0.0244* |
| AF, n (%) | 44 (32) | 23 (31) | 21 (33) | 0.8695 |
| LD, n (%) | 62 (45) | 23 (31) | 21 (33) | 0.2548 |
| CHD, n (%) | 71 (51) | 38 (51) | 33 (52) | 1 |
| CAS, n (%) | 14 (10) | 7 (9) | 7 (11) | 1 |
| HT, n (%) | 5 (4) | 3 (4) | 2 (3) | 1 |
| Therapy, n<br>(rtPA#/APT only) | 33/106 | 20/55 | 13/51 | 0.3968 |

\*P < 0.05; \*\*P < 0.01; \*\*\*P < 0.001

NIHSS- National Institutes of Health Stroke Scale, mRS- modified Rankin Scale, ST- stroke phenotyping (A- atherosclerosis, S- small vessel disease, C- cor, O- other, D- dissection), WBC- white blood count, PLT- platelet, CRP- C reactive protein, AH- arterial hypertension, DM- diabetes mellitus, AF- atrial fibrillation, LD- lipid disorders, CHD-coronary heart disease, CAS- carotid artery stenosis, HT- Hemorrhagic transformation, rtPA- recombinant tissue plasminogen activator, APT- antiplatelet Patients with a higher score on the NIHSS than those obtaining lower scores showed significantly higher concentrations of TNF-alpha, WBC, CRP, NSE, IL-6 and S100B. (Tab. 2)

The differences between other parameters were not statistically significant.

**Table 2.** The comparison of selected parameters between NIHSS and mRS subgroups.

| NIHSS |  |  |  |  |  |  |
| --- | --- | --- | --- | --- | --- | --- |
| Parameter | Test | p-value | FDR corrected p-value | Post-hoc test p-value B-A | Post-hoc test p-value C-A | Post-hoc test p-value C-B |
| WBC | Anova | 0.00496 | 0.01289 | 0.01596 | 0.05830 | 0.28422 |
| CRP | Kruskal | 0.00083 | 0.00361 | 0.00600 | 0.00926 | 0.10253 |
| NSE | Kruskal | 0.02019 | 0.04374 | 0.02093 | 0.37730 | 0.84634 |
| IL-6 | MLE | 0.00001 | 0.00006 | 0.00116 | 0.00023 | 0.00740 |
| S100B | MLE | <0.00001 | <0.00001 | <0.00001 | 0.00028 | 0.05232 |
| TNFalfa | Peto | 0.00218 | 0.00709 | 0.42851 | 0.00864 | 0.03518 |

| mRankin |  |  |  |  |  |  |
| --- | --- | --- | --- | --- | --- | --- |
| Parameter | Test | p-value | FDR corrected p-value | Post-hoc test p-value E-D | Post-hoc test p-value F-D | Post-hoc test p-value F-E |
| WBC | ANOVA | <0.00001 | 0.00002 | 0.00824 | <0.00001 | 0.00627 |
| CRP | Kruskal | <0.00001 | 0.00002 | 0.00054 | <0.00001 | 0.02640 |
| GRN | ANOVA | 0.01411 | 0.03669 | 0.56047 | 0.01129 | 0.02865 |
| IL-6 | MLE.test | 0.00002 | 0.00007 | 0.00925 | 0.00003 | 0.02894 |
| S100B | Peto | 0.00002 | 0.00002 | 0.02357 | 0.00006 | 0.01736 |

WBC- white blood counts, CRP- C reactive protein, NSE- neuron-specific enolasa, IL-6- interleukin 6, S100B- calcium- binding protein B, TNFalfa- tumor necrosis factor alfa
A ≤ 4 points according to NIHSS, B 5-12, C $>$ 12
D ≤ 2 points according to mRS, E = 3, F $>$ 3

Patients with a higher score on the mRS than those obtaining lower scores showed significantly higher concentrations of WBC, CRP, GRN, IL-6, S100B. (Tab. 2) The differences between other parameters were not statistically significant.

Factors with an independent influence on the neurological status on the first day of stroke were: sex, WBC, PLT, CRP, S100B and IL-6 levels. (Tab. 3)

**Table 3.**
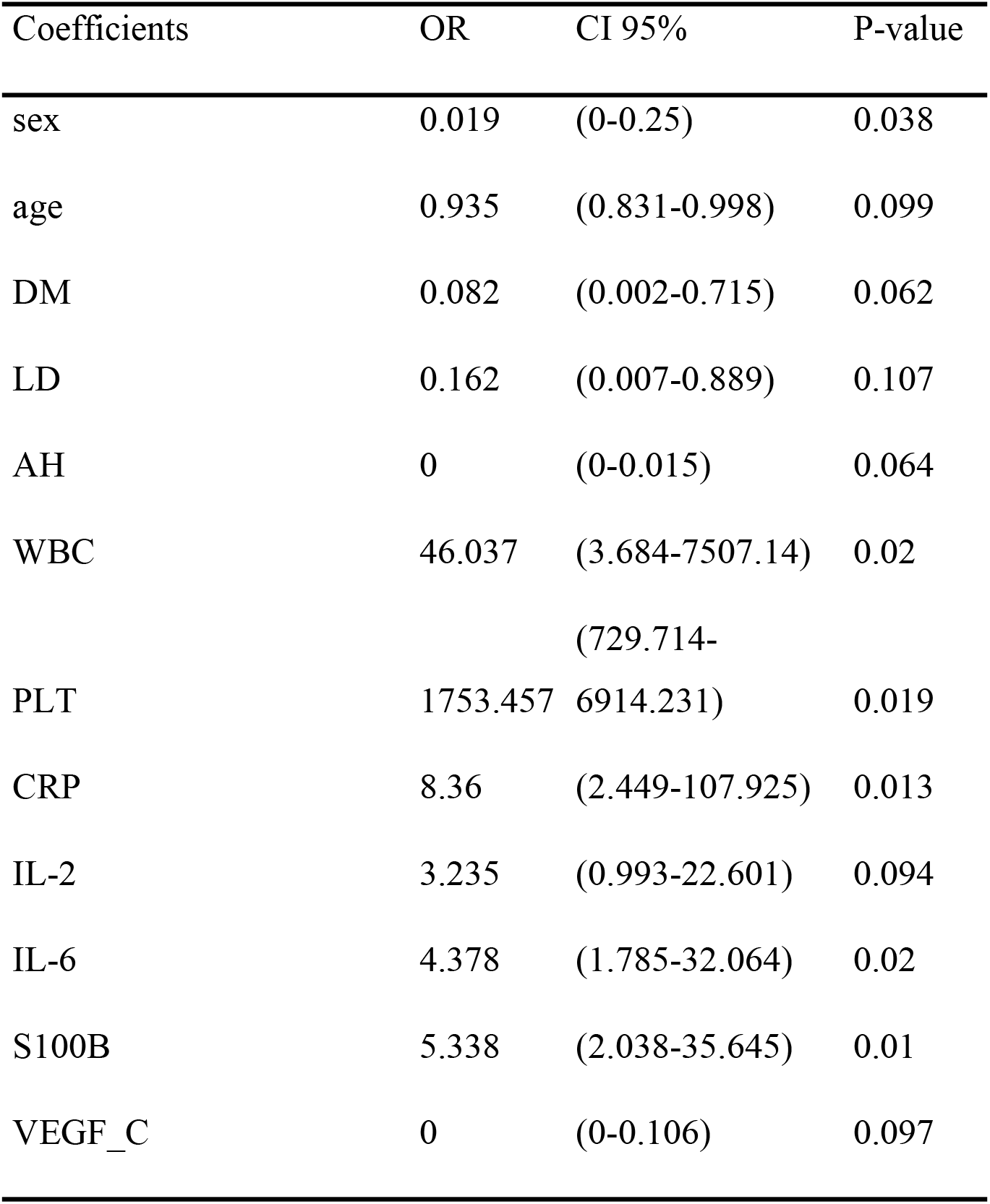
Regression analysis of the influence of clinical phenodata and cytokines on NIHSS

DM- diabetes mellitus LD- lipid disorders AH- arterial hypertension, WBC- white blood count, PLT- platelet, CRP- C reactive protein, IL-6- interleukin 6, IL-2- interleukin 2, S100B- calcium-binding protein B, VEGF-C- vascular endothelial growth factor C

Atrial fibrillation, leukocyte count, CRP, NSA, uPA, interleukin 6 and S100B showed an independent impact on the functional status on the 30th day of stroke. (Tab. 4)

**Table 4.** The regression analysis of the influence of clinical phenodata without gender feature and cytokines on mRS

| Coefficients | OR | CI 95% | P-value |
| --- | --- | --- | --- |
| AF | 6.162 | (1.981-30.917) | 0.006 |
| CHD | 2.75 | (1.065-9.474) | 0.056 |
| CAS | 0.114 | (0.009-0.741) | 0.044 |
| WBC | 23.158 | (3.269-365.421) | 0.005 |
| PLT | 6.22 | (1.166-57.467) | 0.054 |
| CRP | 2.583 | (1.593-5.419) | 0.001 |
| NSE | 0.247 | (0.072-0.573) | 0.004 |
| uPA | 0.102 | (0.008-0.694) | 0.031 |
| IL-6 | 1.512 | (1.069-2.249) | 0.022 |
| S100B | 3.286 | (1.757-8.013) | 0.001 |

mRS- modified Rankin Scale, AF- atrial fibrillation, DM- diabetes mellitus CAS- carotid artery stenosis, WBC- white blood count, PLT- platelet, CRP- C reactive protein, NSE- neuron-specific enolase, uPA- urokinase-type plasminogen activator, IL-6- interleukin 6, S100B- calcium-binding protein B, VEGF-C- vascular endothelial growth factor C

Patients who reached a good functional status on the 30th day, despite an initial poor neurological status, as compared to others, had significantly higher TNF alpha, NSE, IL-2 and VEGF levels. (Tab. 5)

**Table 5.** Logistic regression analysis of the influence of cytokines into the neurological improvement in patients with the best clinical benefits.

| Coefficients | OR | CI 95% | P-value |
| --- | --- | --- | --- |
| BDNF | 22.809 | (1.451-222.998) | 0.098 |
| TNFA | 9.865 | (9.777-10.145) | 0.04 |
| VEGF | 0.695 | (0.685-0.898) | 0.054 |
| NSE | 3049.504 | (13.636-1.58E+04) | 0.044 |
| IL-2 | 0.001 | (0.001-0.123) | 0.044 |
| S100B | 0.002 | (0.001-0.002) | 0.057 |
| VEGF_C | 4.48E+05 | (83.673-1.56E+06) | 0.036 |

BDNF- brain-derived neurotrophic factor, TNFA- tumor necrosis factor alpha NSE- neuron-specific enolase, IL-2- interleukin 2, S100B- calcium-binding protein B, VEGF-C- vascular endothelial growth factor C

Patients with symptomatic atherosclerosis of carotid/cerebral and/or coronary arteries, as compared to others, were older (p=0.003) and had higher levels of CRP, Il-6, and S100B. In each case, the differences were statistically significant (Tab. 6).

**Table 6.** Logistic regression analysis of the importance of cytokines in patients with symptomatic atherosclerosis.

| Coefficients | OR | CI 95% | P-value |
| --- | --- | --- | --- |
| Age |  |  | 0.003 |
| BDNF | 0.705 | (0.365-1.062) | 0.207 |
| WBC | 3.215 | (1.043-13.254) | 0.068 |
| CRP | 0.603 | (0.346-0.91) | 0.025 |
| uPA | 3.653 | (0.904-24.63) | 0.124 |
| IL-6 | 0.593 | (0.324-0.88) | 0.017 |
| S100B | 1.646 | (1.044-3.158) | 0.036 |
| VEGF_C | 0.207 | (0.01-1.16) | 0.106 |

BDNF- brain-derived neurotrophic factor, WBC- white blood count, CRP- C reactive protein, uPA- urokinase-type plasminogen activator, IL-6- interleukin 6, S100B- calcium-binding protein B, VEGF-C- vascular endothelial growth factor C

There were clear correlations between the following factors: a positive correlation for the NSE and S100B pair, and for the BDNF and VEGF, as shown in Figure 1. (Fig. 1)

**Figure 1.**
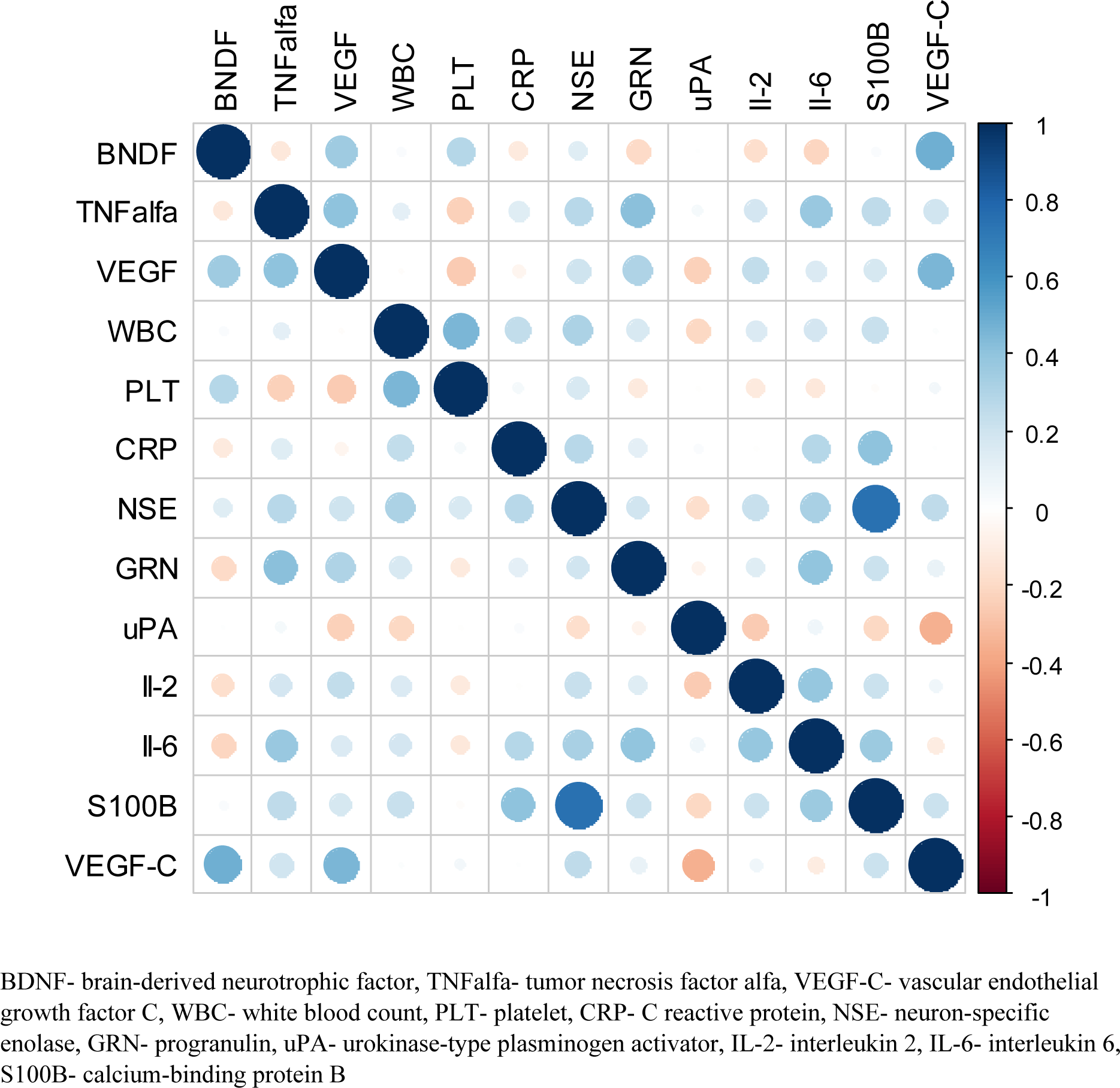
Correlation plot of all analysed features.

## Discussion

Acute cerebral ischemia triggers local and systemic immune response. Immediately after the ischemic event, microglia, mast cells, and astrocytes are activated. These form the initial source of cytokines which cause the BBB permeability and encourage the migration of inflammatory cells from the periphery into the brain. [11] During the first few hours, neutrophils penetrate into the hypoxemic tissue which is invaded by macrophages and monocytes on consecutive days. [12] These cells are known to produce cytokines, free radicals, metalloproteinases, nitric oxide and much more. Such substances directly harm the nerve tissue, induce apoptosis and damage the blood-brain barrier. The destruction to the BBB makes it permeable not only for leukocytes but also for water, which may cause the local edema. [13] Therefore, it has long been stressed that there is a connection between the severity of inflammation, the size of the area affected by ischemia and the degree of neurological deficit. [14] In this study, both the increased leukocyte count and the CRP on the first day of the disease were significantly associated with a more severe neurological status in the ultra-acute phase of the disease and with the functional status on the 30th day. Our results are consistent with previous reports and may be supportive of anti-inflammatory strategies in the treatment of stroke. [15] Studies on experimental stroke suggest that cytokines modulate the process of neuronal damage during ischemic stroke. Their effect on the evolution of infarct depends on their presence and the possibly on their concentration in the area of the penumbra. The interrelationship between ischemia and the activation of the inflammatory process is complex, and the division of acute stroke into stages of destruction and subsequent reconstruction is artificial, as these processes are most likely simultaneous. As cytokines take an active part in the evolution of the cerebral infarction zone, their measurement may serve as a marker of stroke severity in the acute phase of the disease and also as a marker of the outcome in later stages. [16,17] TNF-alpha, IL-1 beta, IL-6, and chemokines are the mediators of an early inflammatory response. [1]

Although TNF and interleukin 1, 2 and 6 are the most frequently studied cytokines in the CSF and blood of stroke patients, the conclusions are ambiguous and inconsistent with the results of experimental stroke studies. [2–4]

This study shows that a high TNF concentration on the first day of stroke correlates with a severe neurological status during that period, while a high IL-6 concentration is not only associated with the severity of neurological status during the early hours of stroke but also with a poor functional status on the 30th day of the disease. IL-6 expression was significantly higher in patients with symptomatic atherosclerosis in the coronary and/or carotid/cerebral arteries. There was no significant correlation between IL-2 and the clinimetric parameters during the first month of the disease. Experimental studies showed that TNF concentration is low after four hours of stroke, with a subsequent increase sustained for several days. [18] It is suggested that TNF has an unfavorable effect on the volume of stroke infarct, in contrast to the effect of IL-6, although there are also reports that contradict these relationships. Therefore, it is possible that the effect depends on the concentration of these cytokines. [19–24] Regardless of individual reports on the neuroprotective potential of TNF and IL-6, just like reported by other investigators, we observed an adverse effect of high concentrations of these cytokines on the neurological status and the degree of post-stroke disability in the first month following onset. [25–28]

NSE and S100B are considered as biomarkers of glial cell damage or neuronal damage in patients with stroke or injury of the CNS. [29–37] Increased blood concentrations of NSE and/or S100B occur after hypoxia, hemorrhage, and injury to the CNS and always indicate that the BBB has been damaged. [38,39] We obtained a positive correlation between the presence of NSE and a worse neurological status on the first day, and between S100B and a poor status on the first and 30th day of stroke, which is consistent with the observations reported by other authors. [15, 40–42] We observed a positive correlation between the concentrations of NSE and S100B.

In recent years, researchers focused on progranulin (GRN), a glycoprotein growth factor with pleiotropic effects on the nervous system. Despite the results suggesting its protective role against ischemic brain injury, little is known about its importance for the prognosis. The results of experimental studies suggest that progranulin prevents BBB damage, causes neuroimmune suppression, is a regulator of vascular permeability, reduces the local brain edema and the size of infarct. In animal studies, it turned out to be a prognostic factor for motor functions. [43–44] In our study, we obtained a positive correlation between GRN concentration on the first day of stroke and the severity of the functional status on the 30th day of the disease. In studies conducted by Xie et al., GRN was a factor related to poor prognosis in post-stroke patients during a 6-month follow-up. [45] Although unfavorable early results were reported in clinical practice, there are experimental data related to the positive effect to reduce the risk of hemorrhagic transformation of an infarct lesion. [46]

The results of this study indicate that the VEGF and BDNF have an effect of summation. VEGF at low concentrations stimulates moderate angiogenesis and prevents delayed neuronal death while reducing the cytotoxic effect of glutamate, thereby increasing cell survival. [47–49] In addition, VEGF has a strong anti-inflammatory effect and promotes neuroplasticity, further enhancing the migration and proliferation of neuronal precursor cells. [50] However, a high concentration of VEGF promotes strong angiogenesis in the hypoxemic area, which can lead to the local edema and to a worsening of the prognosis. [51] Similarly, BDNF increases the secretion of the anti-inflammatory cytokine, has a neuroprotective role in ischemic stroke, and stops inflammatory processes by modifying the MAPK pathway and Bcl-2 cascades. [52, 53]

The results of our study presented here show that the inflammation and damage to the nervous tissue and the BBB dominate in the acute period of cerebral ischemia, which is manifested by the expression of factors being active in the processes mentioned. Based on our observations, it seems that the above issues are important for the neurological status in the early hours of the disease and for the functional status one month after the onset. The expression of inflammatory agents is particularly exhibited in patients with symptomatic atherosclerosis. However, a novelty in our study is the demonstration of the primary role of the inflammatory state, whose parameters show a consistent effect on the degree of neurological deficit. We did not find a significant influence of the selected parameters of the angiogenesis or neurogenesis on the prognosis of patients’ condition in the first month of the disease.

Regardless of the self-limiting nature of inflammation during acute cerebral ischemia, the use of modulators of this process may protect the nervous tissue from permanent damage and improve the prognosis for post-stroke disability.

## Limitations

The limitation of our study is the lack of analysis related to the relationship between the size of an ischemic lesion in head CT/MRI and the blood serum parameters selected for this project.

## Conclusions

The concentration of Il-6 and S100B on the first day of stroke are significant for both the neurological status and the functional status in the acute period of the disease.

Increased CRP and leukocyte count are associated with a worse prognosis regarding the course of acute stroke.

The expression of pro-inflammatory agents and markers of blood-brain barrier damage in the acute phase of stroke is more prominent in patients with symptomatic atherosclerosis than in patients with no clinical features of atherosclerosis.

In the acute phase of stroke, there is a clear, positive correlation between factors of a similar profile of action.

The expression of inflammatory parameters indicates the importance of the inflammatory process, starting during the early days of ischemic stroke, for the post-stroke neurological deficit.

## The Conflict of Interest Declaration

The authors don’t declare the conflict of interest.

## References

1. Chen J, Xiaoming H, Stenzel-Poore M, Zhang J. Immunological mechanism and therapies in brain injures and stroke. Springer; New York 2014.

2. Offner H, Vandenbark AA, Hurn PD. Effect of experimental stroke on peripheral immunity: CNS ischemia induces profound immunosuppression. Neuroscience 2009; 158: 1098–111. doi: 10.1016/j.neuroscience.2008.05.033.

3. Chapman KZ, Dale VQ, Dénes A, et al. A rapid and transient peripheral inflammatory response precedes brain inflammation after experimental stroke. J Cereb Blood Flow Metab 2009; 29: 1764–1768. doi: 10.1038/jcbfm.2009.113.

4. Chang L, Chen Y, Li J, et al. Cocaine-and amphetamine-regulated transcript modulates peripheral immunity and protects against brain injury in experimental stroke. Brain Behav Immun. 2011; 25: 260–269. doi: 10.1016/j.bbi.2010.09.017.

5. Young FB, Weir CJ, Lees KR: GAIN International Trial Steering Committee and Investigators. Comparison of the National Institutes of Health Stroke Scale with disability outcome measures in acute stroke trials. Stroke 2005; 36: 2187–2192.

6. Weisscher N, Vermeulen M, Roos YB, de Haan RJ: What should be defined as good outcome in stroke trials; a modified Rankin score of 0–1 or 0–2? J Neurol 2008; 255: 867–874.

7. Catapano A, Graham I, De Backer G, et al. Guidelines for the Management of Dyslipidaemias. Eur Heart J 2016; 39: 2999–3058.

8. Liakos CI, Grassos CA, Babalis DK, et al. 2013 ESH/ESC guidelines for the management of arterial hypertension: what has changed in daily clinical practice? High Blood Press Cardiovasc Prev 2015; 22: 43–53.

9. Gionfriddo MR, McCoy RG, Lipska KJ. The 2013 American Association of Clinical Endocrinologists’ diabetes mellitus management recommendations: improvements needed. JAMA Intern Med 2014; 174: 179–180.

10. North American Symptomatic Carotid Endarterectomy Trial Collaborators. Beneficial effect of carotid endarterectomy in symptomatic patients with high-grade carotid stenosis. N Engl J Med 1991; 325: 445–453.

11. Lindsberg PJ, Strabian D, Karjalainen-Lindsberg ML. Mast cells as early responders in the regulation of acute blood-brain barier changes after cerebral ischemia and hemorrhage. J.Cereb.Blood.Flow Metab 2010; 30: 689–702.

12. McKittrick CM, Lawrence CE, Carswell H. Mast cells promate blood brain barier breakdown and neutrophil infiltration in a mouse modelof focal cerebral ischemia. J.Cereb.Blood.Flow Metab 2015; 35: 638–647.

13. Nilupul PM, Ma HK, Arawaka S, et al. Inflammation following stroke. J.Clin. Neurosci 2006; 13: 1–8.

14. Kocic I, Kowianski P, Rusiecka I, et al. Neuroprotective effect of masitinib in rats with postischemic strok. Arch Pharmacol 2015; 388: 79–86.

15. Pandey A, Shrivastava AK, Saxena K. Neuron specific enolase and c-reactive protein levels in stroke and its subtypes: correlation with degree of disability. Neurochem Res 2014; 39: 1426–1432. doi: 10.1007/s11064-014-1328-9.

16. Jickling GC, Sharp FR. Blood biomarkers of ischemic stroke. Neurotherapeutics 2011; 8: 349–360.

17. Sharp FR, Jickling GC, Stamova B, et al. Molecular markers and mechanisms of stroke: RNA studies of blood in animals and humans. J Cereb Blood Flow Metab 2011; 31: 1513–1531. doi: 10.1038/jcbfm.2011.45.

18. Lambertsen KL, Clausen BH, Babcock AA, et al. Microglia protect neurons against ischemia by synthesis of tumor necrosis factor. J Neurosci 2009; 29: 1319–1330. doi: 10.1523/JNEUROSCI.5505-08.2009.

19. Taoufik E, Valable S, Müller GJ. et al. FLIP(L) protects neurons against in vivo ischemia and in vitro glucose deprivation-induced cell death. J Neurosci 2007; 27: 6633–6646.

20. Lavine SD, Hofman FM, Zlokovic BV. Circulating antibody against tumor necrosis factor-alpha protects rat brain from reperfusion injury. J Cereb Blood Flow Metab 1998;18;52–58.

21. Yang GY, Gong C, Qin Z, et al. Inhibition of TNF alpha attenuates infarct volume and ICAM-1 expression in ischemic mouse brain. Neuroreport 1998; 9:2131–2134.

22. Herrmann O, Tarabin V, Suzuki S, et al. Regulation of body temperature and neuroprotection by endogenous interleukin-6 in cerebral ischemia. J Cereb Blood Flow Metab 2003; 23:406–415.

23. Clark WM, Rinker LG, Lessov NS, et al. Lack of interleukin-6 expression is not protective against focal central nervous system ischemia. Stroke 2000; 31: 1715–1720.

24. Zaremba J, Losy J. Early TNF-alpha levels correlate with ischaemic stroke severity. Acta Neurol Scand 2001; 104:288–295.

25. Beridze M, Sanikidze T, Shakarishvili R, et al. Selected acute phase CSF factors in ischemic stroke: findings and prognostic value. BMC Neurol 2011; 11: 41. doi: 10.1186/1471-2377-11-41.

26. Waje-Andreassen U, Kråkenes J, Ulvestad E, et al. IL-6: an early marker for outcome in acute ischemic stroke. Acta Neurol Scand 2005; 111: 360–365.

27. Smith C, Emsley H, Gavin C, et al. Peak plasma interleukin-6 and other peripheral markers of inflammation in the first week of ischaemic stroke correlate with brain infarct volume, stroke severity and long-term outcome. BMC Neurol 2004; 4: 2. doi: 10.1186/1471-2377-4-2

28. Mazzotta G, Sarchielli P, Caso V, et al. Different cytokine levels in thrombolysis patients as predictors for clinical outcome. Eur J Neurol 2004; 11: 377–381.

29. Bloomfield SM, McKinney J, Smith L, Brisman J. Reliability of S100B in predicting severity of central nervous system injury. Neurocrit. Care 2007; 6: 121–138.

30. Adami C, Sorci G, Blasi E, et al. S100B expression in and effects on microglia. Glia 2001; 33: 131–142.

31. Casmiro M, Maitan S, De Pasquale F, et al. Cerebrospinal fluid and serum neuron-specific enolase concentrations in a normal population. Eur J Neurol 2005; 12: 369–374.

32. Jauch EC, Lindsell C, Broderick J, et al. Association of serial biochemical markers with acute ischemic stroke: The National Institute of Neurological Disorders and Stroke recombinant tissue plasminogen activator Stroke Study. Stroke 2006; 37: 2508–2513.

33. Bustamante A, López-Cancio E, Pich S, et al. Blood Biomarkers for the Early Diagnosis of Stroke: The Stroke-Chip Study. Stroke; 2017; 48: 2419–2425. doi: 10.1161/STROKEAHA.117.017076.

34. Heidari K, Asadollahi S, Jamshidian M, et al. Prediction of neuropsychological outcome after mild traumatic brain injury using clinical parameters, serum S100B protein and findings on computed tomography. Brain Inj 2015; 29: 33–40.

35. Nylen K, Ost M, Csajbok LZ, et al. Serum levels of S100B, S100A1B and S100BB are all related to outcome after severe traumatic brain injury. Acta Neurochir (Wien) 2008; 150: 221–227.

36. Thelin EP, Nelson DW, Bellander BM. A review of the clinical utility of serum S100B protein levels in the assessment of traumatic brain injury. Acta Neurochir (Wien) 2017; 159: 209–225.

37. Samanci Y, Samanci B, Sahin E, et al. Neuron-specific enolase levels as a marker for possible neuronal damage in idiopathic intracranial hypertension. Acta Neurol Belg 2017; 117: 704–711.

38. Goksuluk H, Gulec S, Ozcan OU, et al. Usefulness of Neuron-Specific Enolase to Detect Silent Neuronal Ischemia After Percutaneous Coronary Intervention. Am J Cardiol 2016; 117:1917–1920. doi: 10.1016/j.amjcard.2016.03.037.

39. Kanavaki A, Spengos K, Moraki M, et al. Serum Levels of S100b and NSE Proteins in Patients with Non-Transfusion-Dependent Thalassemia as Biomarkers of Brain Ischemia and Cerebral Vasculopathy. Int J Mol Sci 2017;18: 12. pii: E2724. doi: 10.3390/ijms18122724.

40. Nayak AR, Badar SR, Lande N, et al. Prediction of Outcome in Diabetic Acute Ischemic Stroke Patients: A Hospital-Based Pilot Study Report. Ann Neurosci. 2016; 23: 199–208.

41. Hu Y, Meng R, Zhang X, et al. Serum neuron specific enolase may be a marker to predict the severity and outcome of cerebral venous thrombosis. J Neurol 2018; 265: 46–51. https://doi.org/10.1007/s00415-017-8659-9

42. Kanazawa M, Takahashi T, Nishizawa M, Shimohata T. Therapeutic Strategies to Attenuate Hemorrhagic Transformation After Tissue Plasminogen Activator Treatment for Acute Ischemic Stroke. J Atheroscler Thromb. 2017; 24: 240–253. doi: 10.5551/jat.RV16006.

43. Jackman K, Kahles T, Lane D, et al. Progranulin Deficiency Promotes Post-Ischemic Blood–Brain Barrier Disruption. J Neurosci 2013; 33: 19579–19589. doi: 10.1523/JNEUROSCI.4318-13.2013.

44. Zhao C, Bateman A. Progranulin protects against the tissue damage of acute ischaemic stroke. Brain 2015; 138: 1770–1773. doi: 10.1093/brain/awv123.

45. Xie S, Lu L, Liu L, et al. Progranulin and short-term outcome in patients with acute ischaemic stroke. Eur J Neurol 2016; 23: 648–655. doi: 10.1111/ene.12920.

46. Kanazawa M, Kawamura K, Takahashi T, et al. Multiple therapeutic effects of progranulin on experimental acute ischaemic stroke. Brain 2015; 138: 1932–1948. doi: 10.1093/brain/awv079.

47. Lee JW, Bae SH, Jeong JW, et al. Hypoxia-inducible factor (HIF-1)a: its protein stability and biological functions. Exp Mol Med, 2004; 36: 1–12.

48. Sun Y, Jin K, Xie L, et al. VEGF-induced neuroprotection, neurogenesis, and angiogenesis after focal cerebral ischemia. J Clin Invest, 2003; 111: 1843–1851.

49. Sun FY, Guo X.: Molecular and cellular mechanisms of neuroprotection by vascular endothelial growth factor. J Neurosci Res 2005; 79: 180–184.

50. Ruiz de Almodovar C, Lambrechts D, Mazzone M, Carmeliet P. Role and therapeutic potential of VEGF in the nervous system. Physiol Rev 2009; 89: 607–648.

51. Manoonkitiwongsa PS, Schultz RL, McCreery DB, et al. Neuroprotection of ischemic brain by vascular endothelial growth factor is critically dependent on proper dosage and may be compromised by angiogenesis. J. Cereb Blood Flow Metab 2004; 24: 693–702.

52. Lasek-Bal A, Jedrzejowska-Szypulka H, Rozycka J, et al. The presence of Tau protein in blood as a potential prognostic factor in stroke patients. J Physiol Pharmacol 2016; 67: 691–696.

53. Chen AI, Li-Jing X, Yu T, Meng M. The neroprotective roles of BDNF in hypoxic ischemic brain injury. Biomed. Reports 2013; 1: 167–176.

